# Optimizing model comparison for enzymatic mechanism analysis

**DOI:** 10.1101/2020.06.01.127993

**Authors:** Ryan Walsh, Philippe Blain

## Abstract

Characterization of enzyme inhibition in drug development is usually limited to a basic analysis with the classical inhibition models or simply the use of IC50 values. However, a better understanding of enzyme physiology and regulation is seen as key to unraveling and treating the processes associated with stubborn disease targets like Alzheimer’s disease. Recently it has been shown that enzyme regulation, through substrate, inhibitor or activator interactions can be modeled using the summation of binding curves. Here we examine the use of the modular equation permutations, that can be produced through binding curve summation, to fit and evaluate the interactions of abietic acid with protein tyrosine phosphatase nonreceptor type 11. This new sort of analysis will allow for improved insight into the physiological role enzymes play and the consequence their modulation may have in disease progression.

## Introduction

Mechanistic analysis of enzyme systems is recognized as key to the understanding of physiology and disease processes (1). Current enzymatic analysis relies heavily on simplified models that work well with high throughput screening (1). This reliance may be due in part to the large number of models and equations which have been developed to describe complicated kinetic interactions. Recently it has been shown that the modulation of enzyme activity either inhibitory or stimulatory can be modeled using binding curves that equate changes in activity to the fractional association of the modifier with the enzyme population (2; 3). This results in distinct catalytic states that can be defined by the molecules associated with the enzyme (Figure 1). By equating changes in activity with binding curves it is possible to establish a modular expansion of the equations defining these binding curves to identify the catalytic states (4) (Figure 2). Here we take that concept one step further by introducing Hill coefficients to the binding curves improving the usefulness of these equations for modeling complex enzymatic mechanisms. This addition simplifies the creation of models of increasing complexity that can then be applied to datasets to evaluate the likelihood of potential interactions. This methodology is applied to the inhibition of protein tyrosine phosphatase nonreceptor type 11 by abietic acid (5). Protein tyrosine phosphatase nonreceptor type 11 is a member of a family of phosphatases that exhibit dynamic substrate interactions, in that they are known to rearrange when interacting with their larger peptide targets (6) and when interacting with small substrate analogues some of them are subject to substrate inhibition (7; 8). As this family of enzymes display complex regulation both by drug candidates (9) and their substrates the ability to evaluate multiple mechanisms clearly demonstrates the utility of this approach.

**Figure 1.**
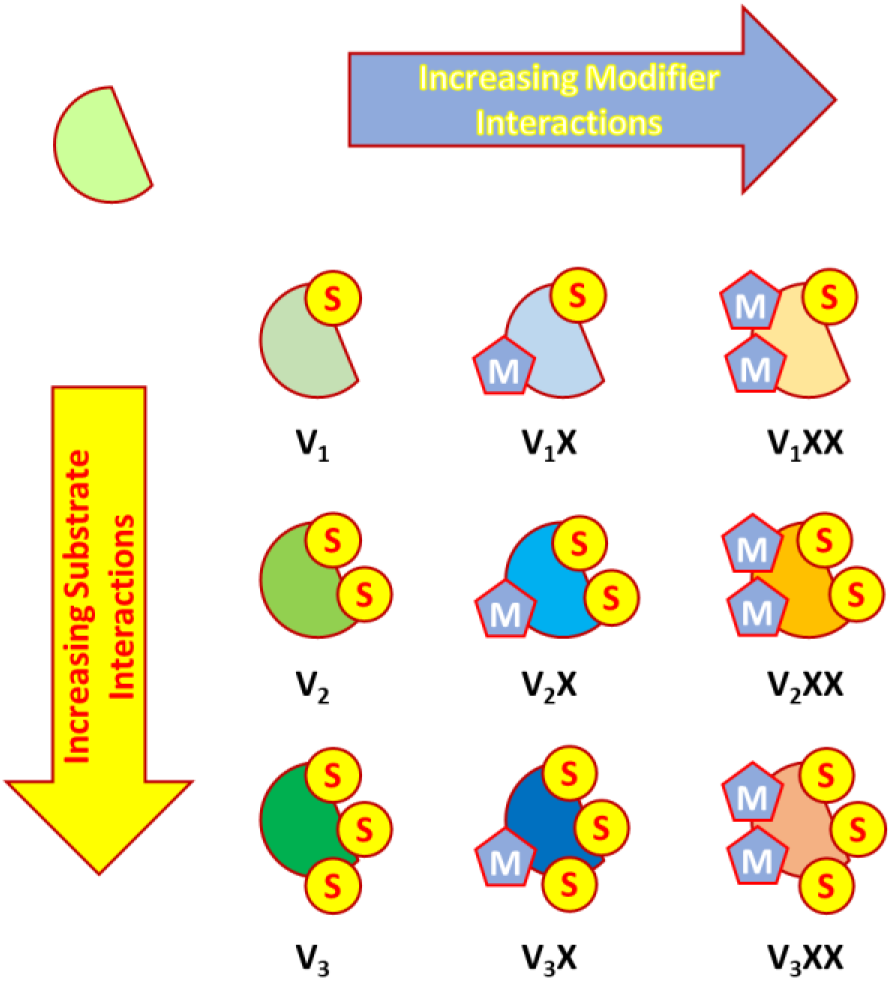
Schematic representing potential catalytically active enzyme states produced by increasing substrate association (S), along with states induced by increasing modifier association (M) which could be either inhibitors or activators.

**Figure 2.**
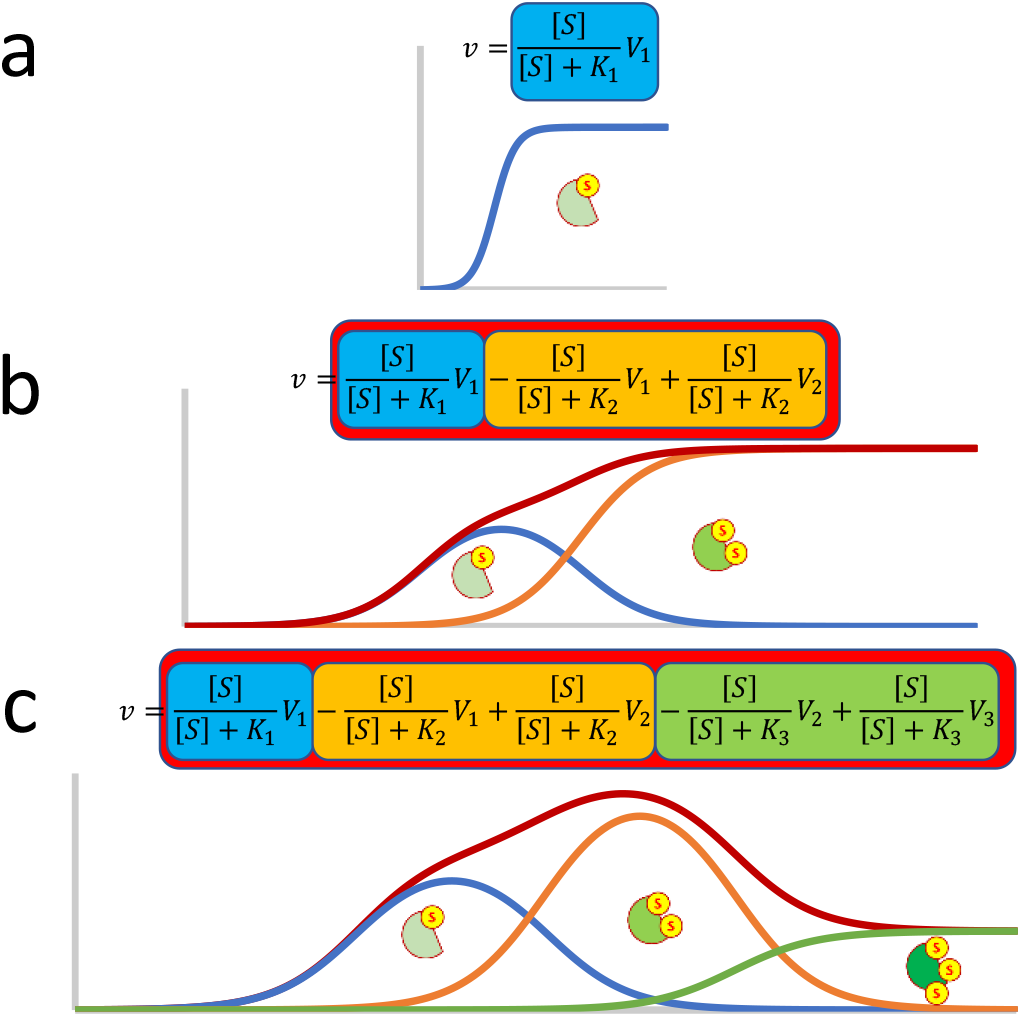
Schematic representing the modular expansion of the (a) Michaelis-Menten equation through the summation of binding curves. In this example (b) secondary binding results in substrate activation (V2 > V1), followed by (c) substrate inhibition (V1 > V3) upon the binding of the third substrate.

## Materials and Methods

The previously published dataset for abietic acid inhibition of protein tyrosine phosphatase nonreceptor type 11 (E.C. 3.1.3.48; accession ID: Q06124) was used in this study (5). All non-linear regression model fitting was done using the solver feature of excel (10). Model development was based upon previously described methodologies (2; 4; 11). Models were evaluated based on their ability to fit the data using the sum of squared residuals (SSR) and Akaike information criterion (AIC) values (12).

## Results and Discussion

In order to analyze the inhibition of SHP2 by AA, ten potential enzyme and inhibitor systems were examined. These systems focused on a maximum of two substrate and two inhibitor interactions with the enzyme (Figure 1; Table 1) resulting in a total of 204 potential inhibition models. As expected, the SSR decreased as the complexity of the models increased. This trend was very apparent with the one substrate one inhibitor system (Table 1a). The addition of Hill coefficients to the substrate-binding curve (Figure 3b; SSR 0.01247) or the inhibitor binding curve (Figure 3c; SSR 0.00899) were clear improvements over the basic equation (Figure 3a; SSR 0.01678). The addition of Hill coefficients to both curves further improved the fit when compared to the other three models (Figure 3d; SSR 0.00420). While in general, increases in model complexity improved the fit, within the two substrate systems a clear improvement in fitting was observed when Hill coefficients were applied to the first substrate-binding term. This trend can be clearly seen in the SSR values associated with the fitting of models in section h where a green streak in column two is associated with the inclusion of a Hill coefficient in the first substrate-binding term but is absent in the third column which lists fits associated with Hill coefficients being added to the second substrate-binding term (Table 1h). This was not unexpected as it is known that SHP2 undergoes a rearrangement when interacting with peptide substrates (6; 13) but appears to experience substrate activation when examined with the small substrate analogue p-nitrophenyl phosphate used in this data set (11).

**Table 1.**
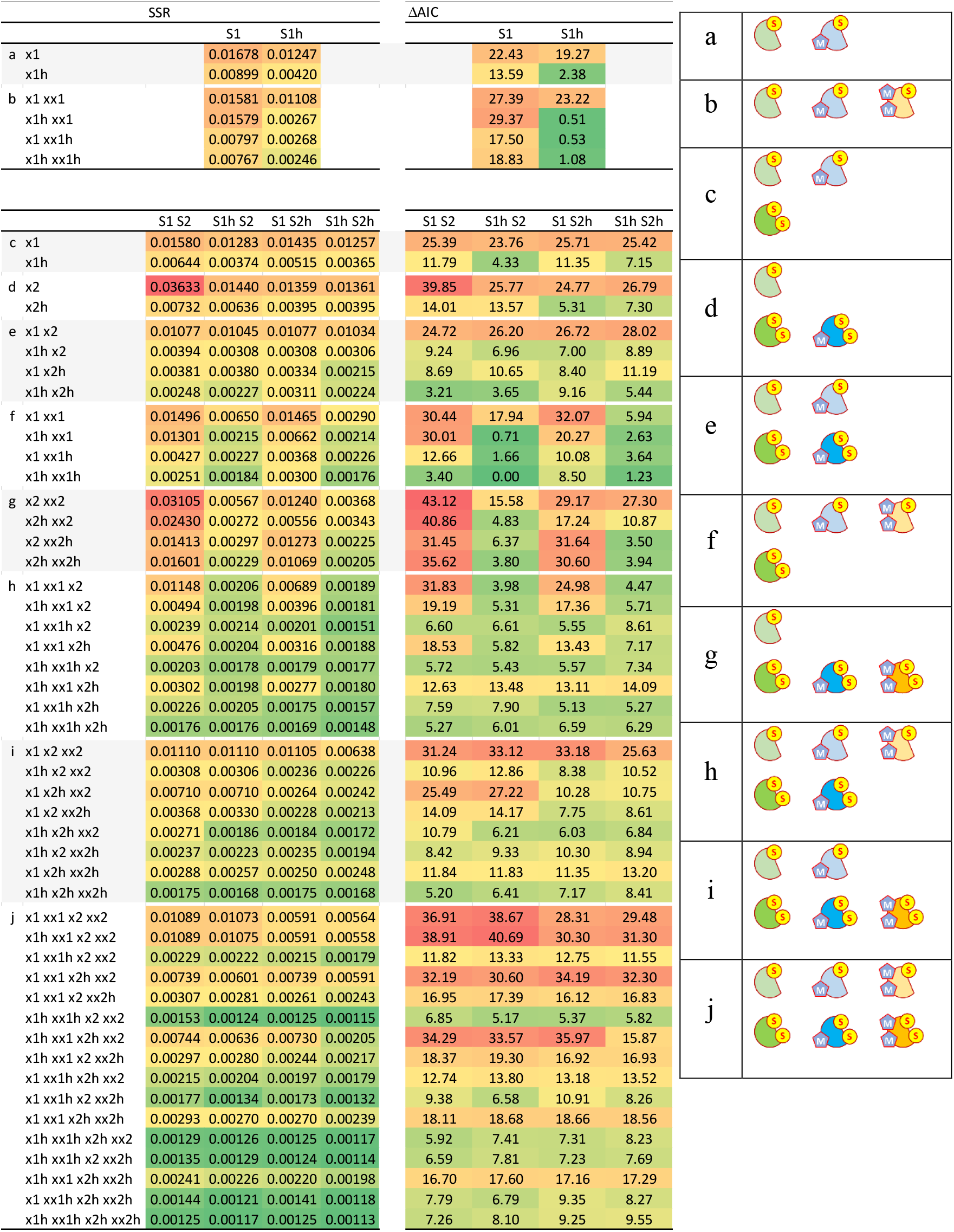
Sum of squared residuals and AIC values associated with the models fit to the SHP2 AA inhibition data. The table is divides based on the inclusion of substrate and modifier induced enzyme states in the model. The columns are defined by substrate-induced forms where S1 indicates one substrate interaction, S2 indicates two and an h indicates the inclusion of Hill coefficients. Rows are defined by potential inhibitor interactions where the number of x’s indicated the number of binding interactions, the number indicates the substrate-induced enzyme form the inhibitor is binding and an h indicates the inclusion of a Hill coefficient. For example, xx2h indicates a second inhibitor binding to the substrate activated enzyme where a Hill modifier is included. The color code is used to highlight better fits (green) from poorer fits (red).

**Figure 3.**
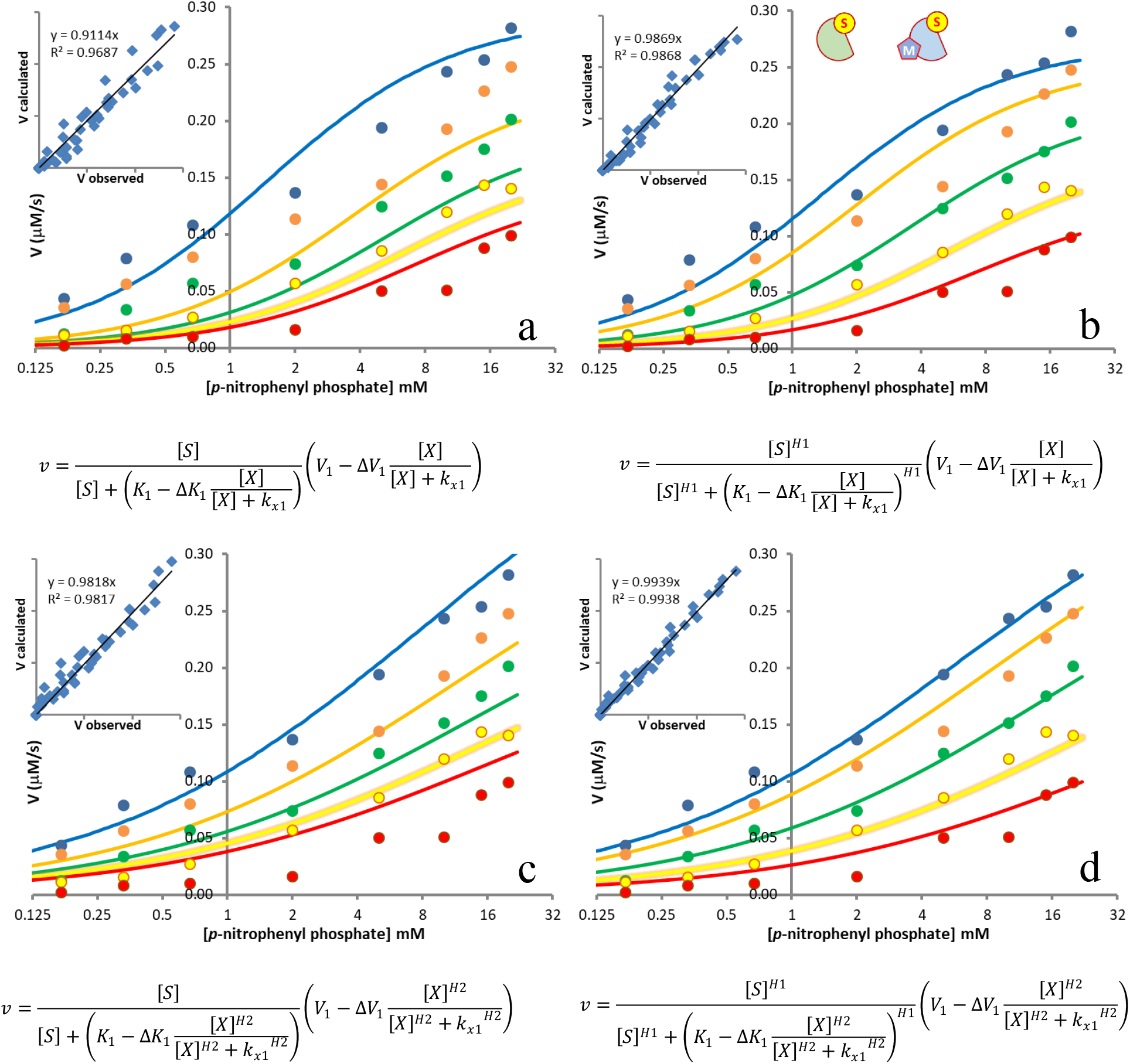
Fitting the SHP2 inhibition by AA dataset to a one substrate one modifier model results in four possible fits, (a) a fit where the Hill coefficient is not applied, (b) a fit where the Hill coefficient is applied to the binding of the substrate, (c) a fit where the Hill coefficient is applied to the binding of the inhibitor and (d) fitting of the model with Hill coefficients applied to both binding interactions. The data set represents the substrate range between 17 μM to 20 mM p-nitrophenyl phosphate with 0 μM (blue), 100 μM (orange), 200 μM (green), 300 μM (yellow) and 400 μM (red) AA. Fittings are accompanied by correlation plots of the calculated versus the observed velocities.

As the complexity of the models ranged between 5 and 22 parameters AIC values were calculated (Table 1) to discriminate between potentially useful models and those which were suffering from overfitting (12). The AIC values suggested that the single substrate model with two Hill coefficients (Figure 3d) potentially provided a useful model of the data (Table 1a). It also indicated that the addition of a second inhibitor to the single substrate system may provide a decent description of the data, with AIC values approaching the global minimum of the data set (Table 1b).

However, it has already been shown that based on plots of the noninhibited enzyme it is probably more appropriate to model the activity of this enzyme using a two-substrate binding model (11). Examining the two substrate models suggested several potentially useful models (Figure 4) associated with single inhibitor interactions with the substrate-induced forms of the enzyme (Table 1c-e). The AIC global minimum appeared with the models describing two inhibitors binding to the non-substrate activated form of the enzyme (Table 1f; Figure 4c). While more complex models improved on the fitting based on SSR values few produced viable models based on their increase in complexity (Figure 4d). Examining the actual binding constants for the model with the AIC minimum suggested a problem with the model, as for this model to achieve the fit the secondary substrate binding constant had inflated to 1.7 M with an associated reaction rate 5.5 μM/s. Both of these values were quite high when contrasted to the substrate range used in the experiments which do not exceed 20 mM and the observed rates 0.28 μM/s. This large increase in rate can be observed as a slightly more pronounced upward swoop in the curve at higher substrate concentrations on the right side of the plot (Figure 4c). The same problem was observed with the simpler single inhibitor model (figure 4a), suggesting this effect was associated with models lacking inhibitor interactions with the substrate activated form of the enzyme. When models were allowed to include inhibitor interactions with the substrate activated form of the enzyme the substrate-binding term for the secondary binding were observed to fall much closer to or within the experimental dataset and changes in catalytic rate were generally doubled rather than suggesting up to twenty-fold increases. Models including inhibition of the substrate activated form also did not display as significant of an upward swoop when compared to models lacking this feature (Figure 4b, 4d). This suggests that while the AIC values of this dataset may favor models where the inhibitor only affects the non-substrate activated form of the enzyme, more appropriate models probably need to include inhibition of the substrate activated form of the enzyme.

**Figure 4.**
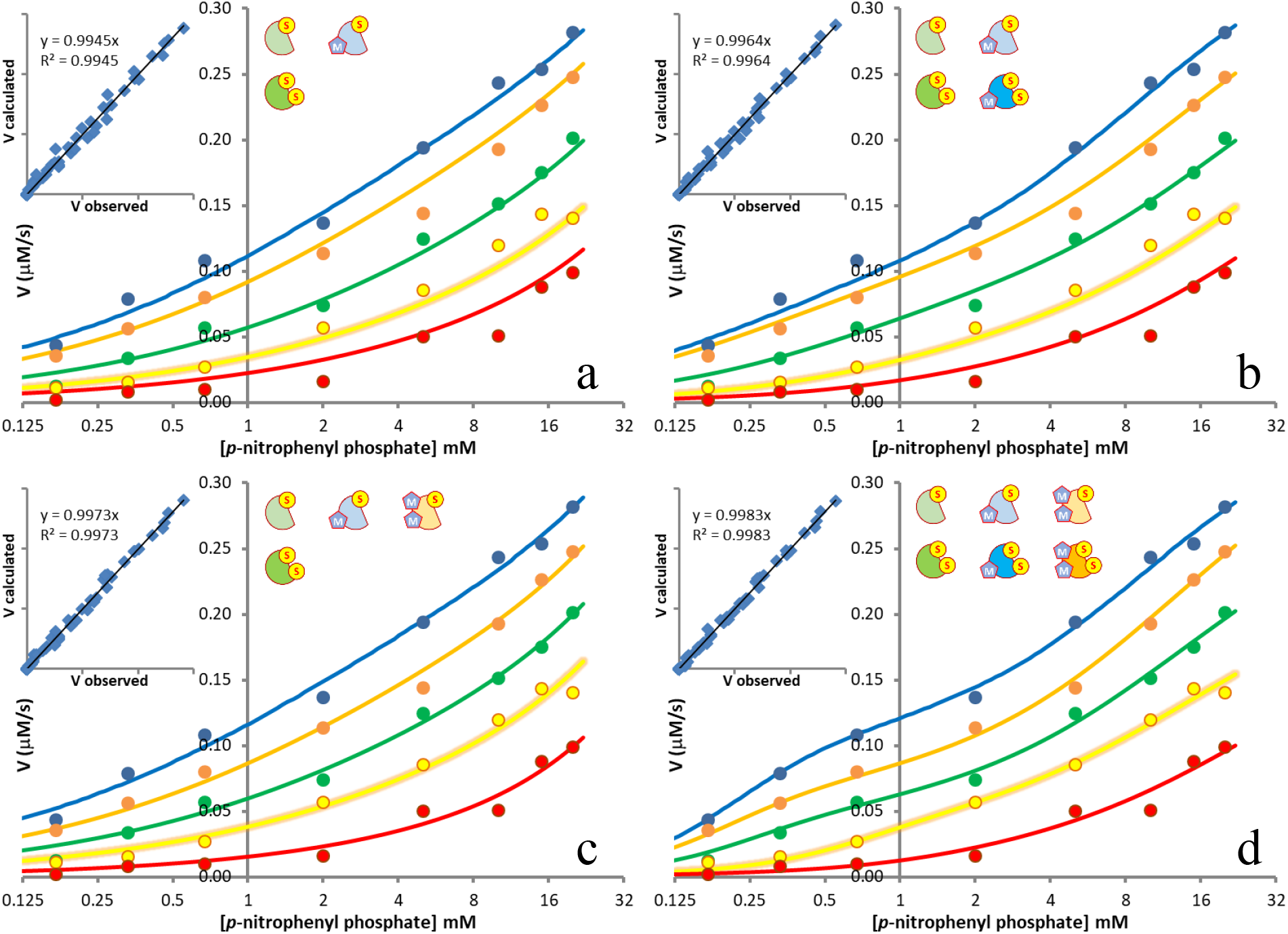
Fitting the of the SHP2 inhibition by AA dataset to the best fit models as defined by the minimum AIC value for the model where (a) only the non-substrate activated enzyme interacts with the inhibitor (b) the inhibitor interacts with both substrate hydrolyzing forms of the enzyme the minimum AIC of the examined models and (d) the minimum sum of squared residuals of the examined models. The data set represents the substrate range between 17 μM to 20 mM p-nitrophenyl phosphate with 0 μM (blue), 100 μM (orange), 200 μM (green), 300 μM (yellow) and 400 μM (red) AA. Fittings are accompanied by correlation plots of the calculated versus the observed velocities.

## Conclusions

The depth and quantity of information provided by this sort of analysis is currently unavailable in drug and biological studies. While currently labor-intensive, the modular and systematic layout of the workflow involved in this study suggests that it will be amenable to automation and integration into modern bioinformatics approaches. Also, the ability to modularly construct complex enzyme kinetic equations should greatly benefit research into disorders such as Alzheimer’s disease where major drug targets like cholinesterases (14) and g-secretase (15; 16) are dynamically regulated by their substrates producing complex interactions with potential drug candidates.

## AUTHOR INFORMATION

## Author Contributions

The manuscript was written through contributions of all authors. All authors have given approval to the final version of the manuscript.

## Notes

## ABBREVIATIONS

AA: abietic acid
ΔAIC: change in Akaike information criterion
SHP2: protein tyrosine phosphatase nonreceptor type 11
SSR: sum of squared residuals

---

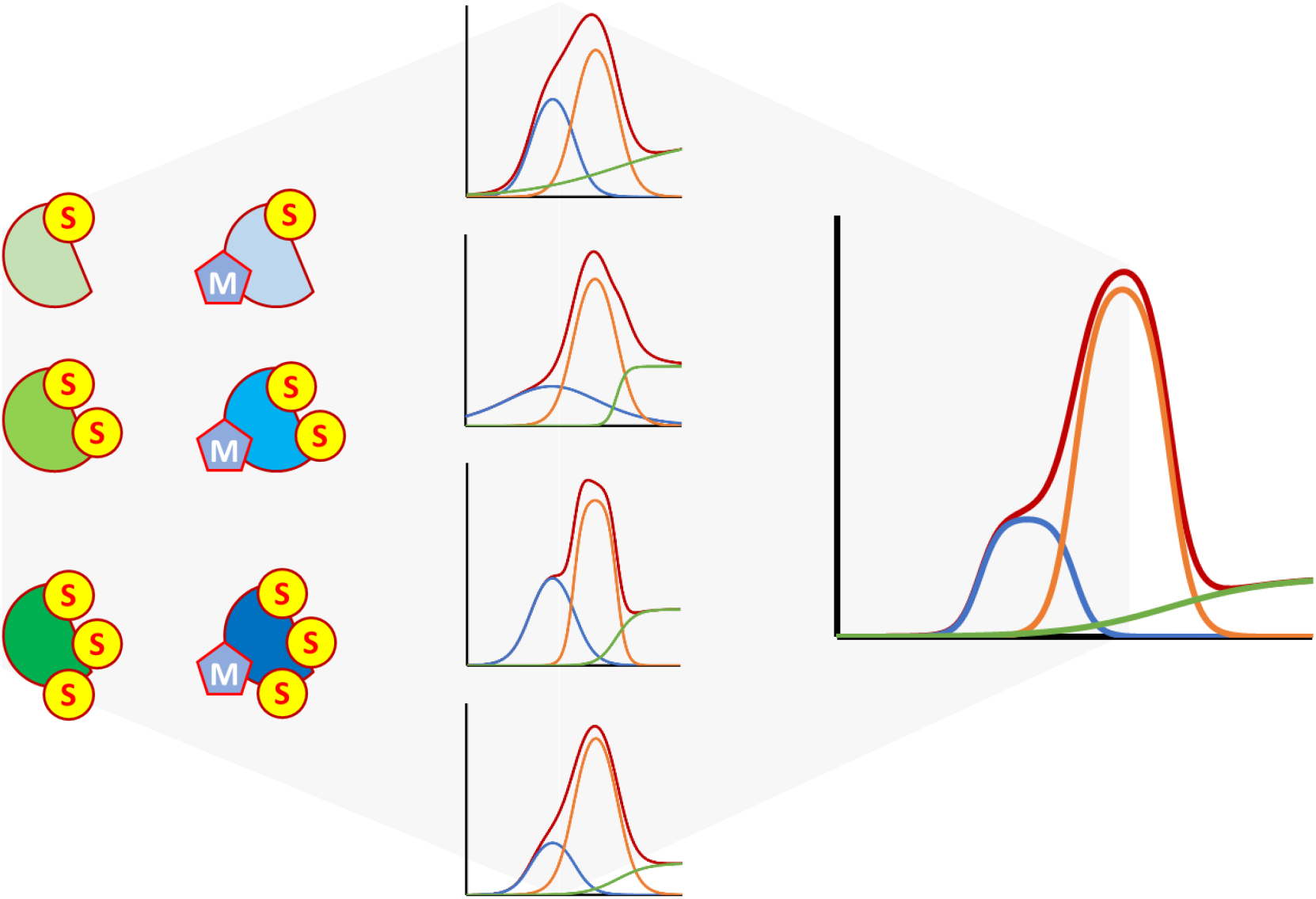

